# Towards a complete diploid chicken genome

**DOI:** 10.64898/2026.10.25.761615

**Authors:** Shiyu Fan, Jiao Liu, Na Kang, Diyan Li, Zhuanjian Li, Naiyi Xu, Luohao Xu

## Abstract

The repetitive and heterochromatic parts of microchromosomes and dot chromosomes have impeded the creation of a complete, haplotype-resolved chicken genome. To overcome these challenges, we employed a trio-sequencing approach, utilizing long reads from a female chick, supplemented by parental long-read data for accurate haplotype phasing. This strategy produced nearly complete, haplotype-resolved paternal and maternal assemblies, with 14 residual gaps. Telomeric sequences remain missing in two paternal and four maternal chromosomes, suggesting a still incomplete assembly. GGswu2 resolved more than 50 Mb of previously unresolved regions, predominantly within dot chromosomes. The dot chromosomes exhibit low nucleotide diversity but a high density of structural variants between haplotypes. We found that the centromere sizes and relative positions of centromere cores sometimes differ between the two haplotype genomes. These findings suggest that comparative genomics must incorporate the high-resolution of these dynamic complex regions to fully understand avian genome evolution and genetic complexity.

## Introduction

The development of highly contiguous and complete vertebrate reference genomes is foundational for understanding evolutionary processes and genetic mechanisms underlying complex traits. The chicken (*Gallus gallus*) is a critically important model organism and agricultural species, yet efforts to fully resolve its genome sequence have long been hampered by its distinctive karyotype^1–4^. The chicken karyotype (2n = 78), like many other bird genomes^5,6^, is made up of 39 chromosome pairs, typically recognized as 10 larger macrochromosomes (GGA1-10) and 28 smaller microchromosomes (GGA11-38), plus the Z and W sex chromosomes^7,8^. The first draft chicken genome, published in 2004, lacked sequence for six chromosomes and suffered from incomplete assembly of the sex chromosomes^9^. Subsequent assemblies continued to face persistent challenges, retaining thousands of gaps and failing to assemble or assign several microchromosomes^10,11^, with a number conserved vertebrate genes missing (“hidden avian genes”)^11–13^. These limitations have prevented a complete view of the avian gene repertoire and genome structure.

Recent advancements in long-read sequencing technologies, particularly PacBio HiFi and Oxford Nanopore Technology (ONT) ultralong sequencing, have enabled the production of complete, telomere-to-telomere (T2T) assemblies for many complex genomes, including the human genome^14–20^. This technological progress facilitated the assembly of GGswu1, published in 2023^21^, which represents the most complete chicken genome assembly to date. Compared with earlier versions such as GRCg7b, GGswu1 introduced a special category of chicken microchromosomes called dot chromosomes (GGA16, 29-32, and 34-38), defined on the basis of epigenetic modifications and genomic repeat content^22–24^. The unique features of dot chromosomes have been reported in many divergent bird lineages^25^, including golden pheasant^26^ and zebra finch^18^, suggesting an ancestral origin of dot chromosomes in birds. Nevertheless, even the GGswu1 benchmark assembly remains technically incomplete, as it still contains unresolved gaps within the highly complex W chromosome, and the short arms of several acrocentric chromosomes are likely incomplete^13^. Moreover, GGswu1 was generated as a compressed haploid sequence, limiting the ability to study allele-specific functions and complex structural variation in a diploid context^17,18,27^.

To address these challenges and achieve a complete diploid (haplotype-resolved) chicken genome, our study employed a trio-based sequencing approach^28^. A female chick, derived from a Daninghe paternal line and Nanchuan maternal line, was sequenced to high coverage using PacBio HiFi, ONT ultralong and Pore-C technologies. Critically, the two parents of the chick were also sequenced using ONT reads. This approach allowed us to leverage the trio-binning strategy for producing highly contiguous haplotype-resolved paternal (GGswu2-p) and maternal (GGswu2-m) assemblies, substantially improving upon GGswu1, particularly in previously unresolved regions of the dot chromosomes, and enabling the analyses presented here.

## Results

### The nearly complete haplotype-resolved genome assembly

We employed a female chick to build a genome with HiFi reads (87 ×), Nanopore reads (79 ×) and Pore-C reads (124 ×). Nanopore reads longer than 100 kb account for 26.8% of the total amount (Table S1). Because some complex regions of chicken microchromosomes are inaccessible for short reads^21^, ∼50 Gb Nanopore reads (along with ∼150 Gb short reads) were also collected for each of the two parents, facilitating trio-based haplotype phasing for microchromosomes.

We employed a hybrid strategy for genome assembly. First, the primary diploid assembly was generated using HiFiasm with the trio-binning mode, using both HiFi and Nanopore ultra-long reads^29^. The two haplotype assemblies have contig N50 values of 37.8 Mb and 36.4 Mb, respectively, both harboring less than 100 gaps. Then we applied the HERRO algorithm to correct ONT reads for genome assembly, and the resulting haplotype-resolved contigs, instead of raw reads, were used for filling gaps of the corresponding haplotype assemblies. The filled regions were systematically evaluated using ONT and HiFi read coverage and read-support analyses, including full-length spanning reads and continuous read support across the filled sequences (Table S2-S3; Fig. S1). We note that gap filling in structurally complex regions may be more susceptible to assembly artifacts and should therefore be interpreted with caution. Overall, the paternal (GGswu2-p) and maternal (GGswu2-m) haplotypes are approximately 1.10 Gb and 1.05 Gb in size, respectively. Only 14 gaps remained in the diploid genome, most of which were located on the dot and W chromosomes (Fig. 1A; Table 1; Table S4).

**Fig. 1.**
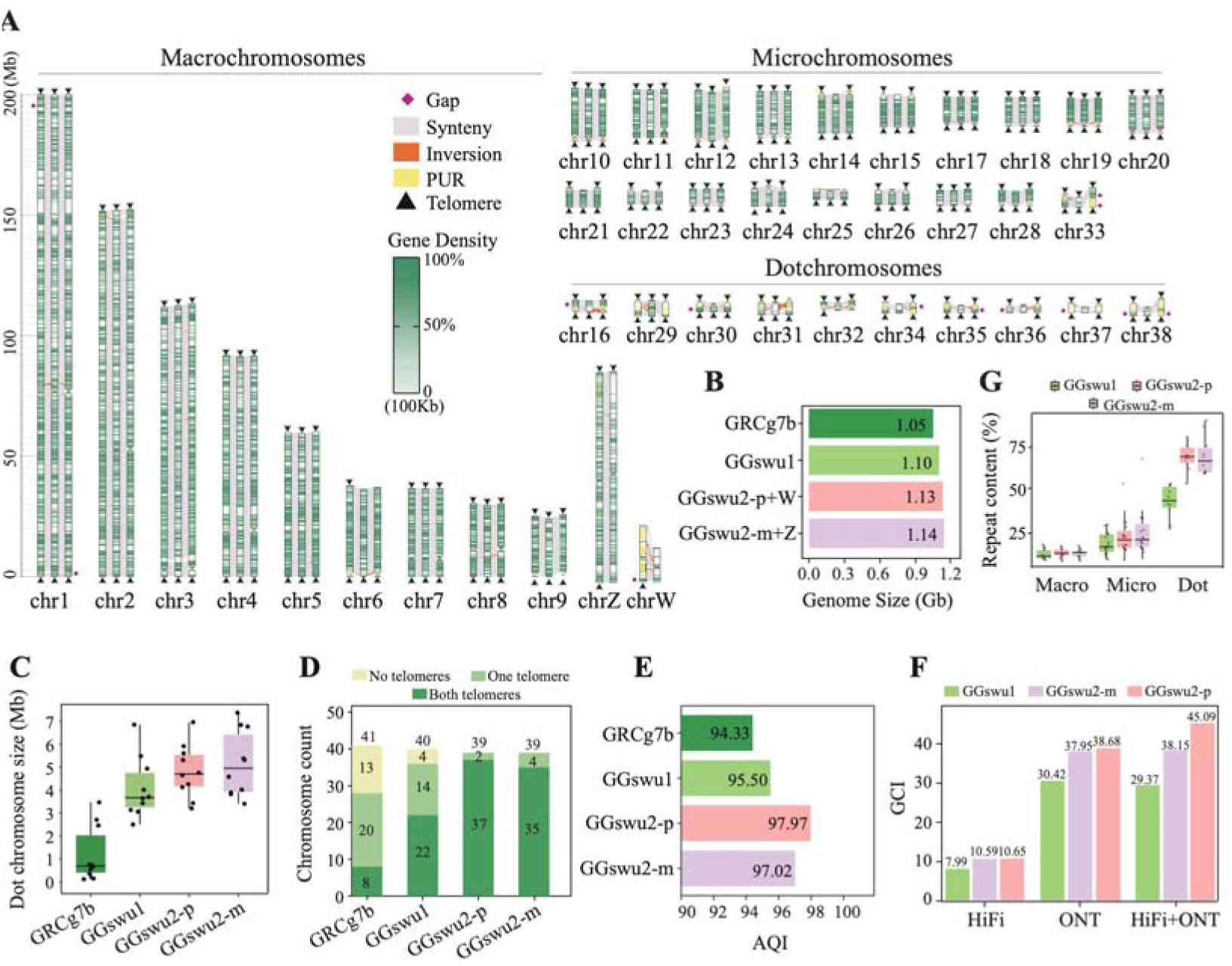
A nearly complete haplotype-resolved chicken genome assembly. (**A**) Chromosomal synteny between GGswu2 and GGswu1. Left: GGswu2-p; middle: GGswu1; right: GGswu2-m. The yellow bars represent the previously unresolved regions (PURs) in GGswu1. The gene density is calculated in 100-kb windows. (**B**) Increased assembly size of GGswu2-p+W and GGswu2-m+Z compared with previous assemblies. The W chromosome was added to GGswu2-p, and the Z chromosome was added to GGswu2-m. (**C**) Increased assembly sizes of dot chromosomes in GGswu2 compared with previous assemblies. (**D**) Summary of telomere completeness across genome assemblies. (**E**) Higher AQI in the GGswu2 assemblies. (**F**) Higher GCI in the GGswu2 assemblies. (**G**) Increased resolution of repetitive sequences in the GGswu2 assemblies.

**Table 1.** Summary of GGswu2 assemblies.

| Statistics | GGswu1 | GGswu2-p | GGswu2-m |
| --- | --- | --- | --- |
| Total length (bp) | 1,100,911,310 | 1,103,386,578 | 1,049,511,662 |
| # chromosomes | 40 | 39 (without W) | 39 (without Z) |
| # gaps | 14 | 6 | 8 |
| # contigs | 54 | 45 | 47 |
| Contig N50 (bp) | 87,735,852 | 91,766,251 | 91,575,483 |
| GC content (%) | 48.52 | 42.63 | 42.87 |
| BUSCOs (%) | 97.0 | 99.2 (GGswu2-p + W) | 99.3 (GGswu2-m + Z) |
| QV | 37.2099 | 48.3711 | 48.1149 |
| GCI (ONT) | 30.4181 | 38.6787 | 37.9486 |
| GCI (HiFi) | 7.9901 | 10.6505 | 10.5924 |
| GCI (ONT+HiFi) | 29.3659 | 45.0861 | 38.1478 |
| AQI ((R-AQI+S-AQI)/2) | 95.50 | 97.97 | 97.02 |

### Comparison with previous genome assemblies

For the sake of direct comparison with previous assemblies, we generated sex-chromosome-complemented assemblies by adding W to GGswu2-p and Z to GGswu2-m, referred to as GGswu2-p+W and GGswu2-m+Z, respectively. Their genome sizes became approximately 1.13 Gb and 1.14 Gb, respectively, larger than those of GGswu1 (1.10 Gb) and GRCg7b (1.05 Gb) (Fig. 1B). The average size of dot chromosomes in GGswu2 was approximately 4.98 Mb, considerably larger than those in GGswu1 (4.07 Mb) and GRCg7b (1.20 Mb) (Fig. 1C). Given that GGswu1 is suggested to be more complete than GRCg7b^13^, we therefore focused on comparisons between GGswu2 and GGswu1 in most of the following analyses.

The performance of the GGswu2 assembly was superior to that of GGswu1 across various analyses. Firstly, GGswu2 has 37 (GGswu2-p) or 35 (GGswu2-m) chromosomes with telomeres at both ends, in contrast to only 22 such chromosomes in GGswu1 (Fig. 1D). Next, by aligning re-sequencing data of different chicken breeds, we showed that the mapping rates for GGswu2-p and GGswu2-m reached 98.14% and 98.09%, respectively, higher than those of GGswu1 (97.94%) or GRCg7b (97.46%, Fig. S2A). The Benchmarking Universal Single-Copy Orthologs (BUSCOs) analyses showed 99.2% completeness for GGswu2-p+W and 99.3% for GGswu2-m+Z, both higher than those of GRCg7b (95.4%) and GGswu1 (97.0%)^13^ (Table 1). To assess base-resolution assembly accuracy, we used Clipping information for Revealing Assembly Quality (CRAQ) that relies on clipping information from read alignments to identify potential errors^30^. We obtained the Assembly Quality Index (AQI) of 97.97 for GGswu2-p and 97.02 for GGswu2-m, again higher than those of GRCg7b (94.33) and GGswu1 (95.5) (Fig. 1E; Table 1). Genome Continuity Inspector (GCI) is another evaluator that quantifies the overall continuity of a genome assembly, particularly targeting assemblies at or near T2T level^31^. We obtained the GCI of 45.09 and 38.15 for GGswu2-p and GGswu2-m based on the combination data of HiFi and ONT, much higher than 29.37 of GGswu1 (Table 1; Fig. 1F).

By aligning GGswu2 against GGswu1, ∼51 Mb previously unresolved regions (PURs) were identified in the GGswu2-p and ∼69 Mb in the GGswu2-m (Fig. 1A; Table S5). We found more PURs in the dot chromosomes (22-26 Mb) than in microchromosomes (12-19 Mb) and macrochromosomes (11-12 Mb). The PURs from the dot chromosomes accounted for ∼50% (both for the two haplotypes) of the total size of GGswu2 dot chromosomes, but only ∼1.4% (GGswu2-p) and 1.6% (GGswu2-m) for the macrochromosomes (Table S5). For instance, we identified 1.9 and 2.1 Mb PURs in the chr35 which had a respective size of only 4.3 (GGswu2-p) and 3.8 Mb (GGswu2-m) (Fig. 1A). In micro- and macrochromosomes, the PURs are mainly located in the subtelomeric and pericentromeric regions (Fig. 1A).

Our GGswu2 assembly also recovered 777 out of the 782 genes that are known to be missing from GGswu1^13^. Those re-discovered genes were mainly located in the dot chromosomes (Fig. S2B). Our analysis therefore suggests that the previous missing genes were mostly in PURs, instead of being the so-called “non-reference sequences”^13^.

Furthermore, despite a substantial increase in the assembled sequences of the dot chromosomes, their genomic and epigenetic features remain unchanged, distinct from that of micro- and macrochromosomes (Fig. S3; Table S6). In fact, because more repetitive sequences have been resolved, the feature of higher repeat content in dot chromosomes becomes more prominent (Fig. 1G).

We next examined whether the short arms of acrocentric chromosomes have been improved in GGswu2 relative to GGswu1. Our analyses showed that the short arms of 30 acrocentric chromosomes have been completed in GGswu2. Those newly resolved sequences have a size of approximately 38 and 40 Mb, accounting for 68.6% and 74.5% of the total PURs of the GGswu2-p and GGswu2-m, respectively (Table S7).

### Resolution of centromeric architecture

Though previous studies have revealed the origin and structure of chicken centromeres^21,32,33^, the variation in centromeric organization has not been fully assessed, in part due to incomplete assembly of the short arms of several acrocentric chromosomes. For instance, the Centromere Protein A (CENP-A) signals were absent in the chromosome 15 and 37 of GGswu1^21^. To test whether this is due to the missing centromeric sequences in the GGswu1 assembly, we mapped the ChIP-seq data of CENP-A against the GGswu2 assemblies. Our analyses recovered the CENP-A peaks on chromosome 15 and 37 of both GGswu2-p and GGswu2-m (Fig. 2A; Fig. S4). In fact, the GGswu2 assemblies resolve the complete sequences of the short-arms of these two acrocentric chromosomes. Similar to many other chicken chromosomes, the CENP-A binding regions of chromosome 15 and 37 consist largely of (TTAGGG)n repeats that are commonly present in the telomeres. The CENP-A flanking regions contain tandem arrays of the CNM repeat (Fig. 2A; Fig. S4).

**Fig. 2.**
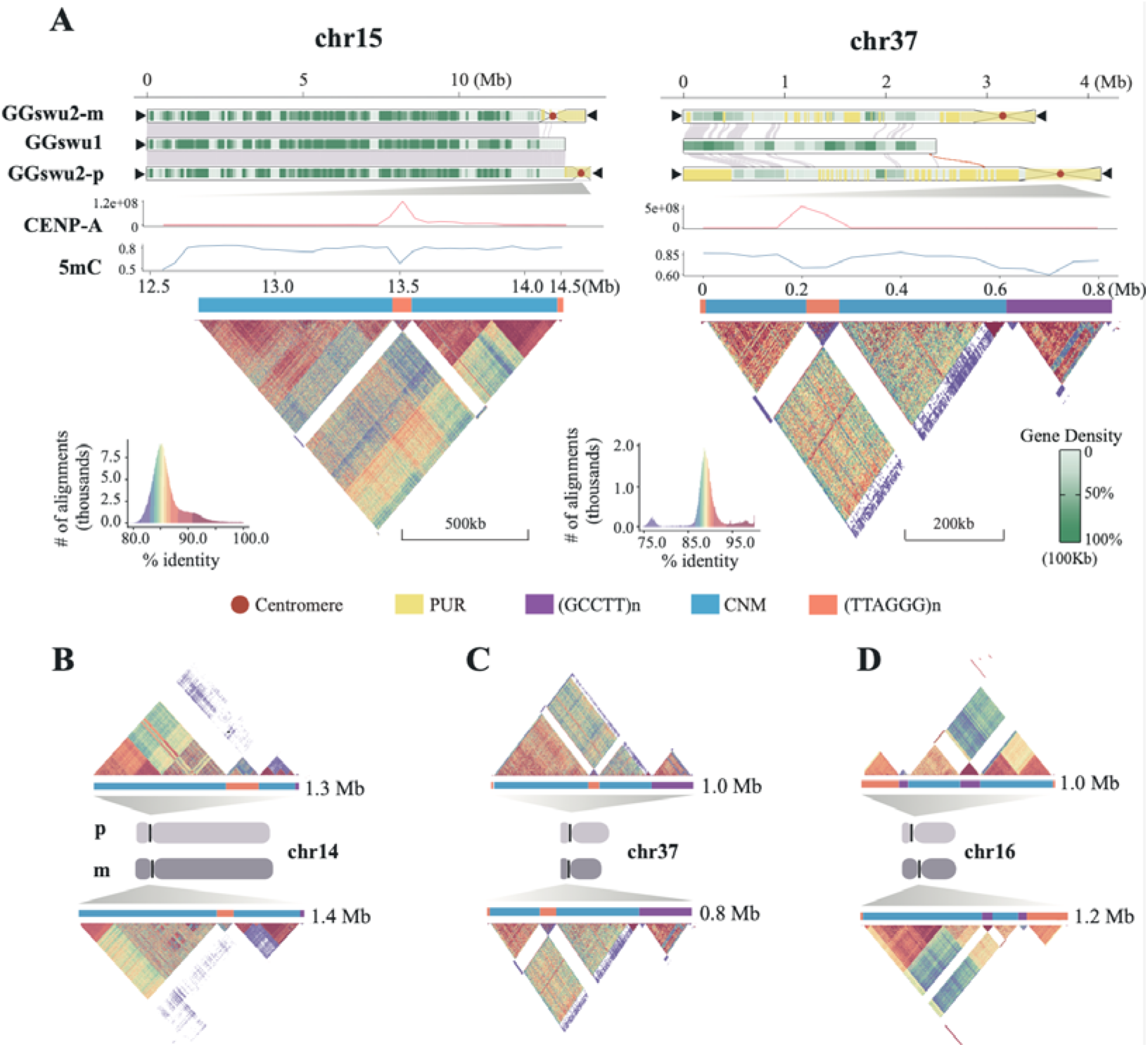
Centromeric architecture and repeat organization. **(A)** Centromeric features and epigenetic landscape in chromosome 15 and 37 (GGswu2-m). Their sequences were missing in GGswu1 because of incomplete genome assembly of the short-arms. The StainedGlass plots for GGswu2-p are shown in the supplementary Figure 4. (**B**) Variation in centromeric organization between two haplotypes of chromosome 14. **(C)** Variation in centromeric organization between two haplotypes of chromosome 37. (**D**) Variation in centromeric organization between two haplotypes of chromosome 16. p: paternal assembly; m: maternal assembly. Inter-haplotypic comparison of other chromosomes are shown in the supplementary Figure 5.

Furthermore, our re-analyses correct several previously misidentified or missing centromeric sequences. For instance, although the CENP-A-occupied centromere core on chromosome 27 was identified in GGswu1, no satellite repeats were detected. In contrast, we identified 0.5 Mb and 0.7 Mb satellite repeats within the centromeres of chromosome 27, respectively (Fig. S4-S5). Moreover, whereas no CNM repeats were reported on chromosome 33 in GGswu1, we detected an approximately 1 Mb CNM repeat array flanked by telomeric repeats in GGswu2 (Fig. S4-S5). On chromosome 38 of GGswu1, multiple atypical CENP-A peaks were detected, but our analyses suggest that the authentic centromere sequence was absent in GGswu1, and we identified distinct and well-resolved centromeric regions on both chromosome 38 haplotypes, spanning approximately 0.74 Mb and 0.80 Mb, respectively (Fig. S4-S5). Although residual gaps remain on chromosome 38, these gaps are located outside the identified centromeric regions and therefore do not affect the resolution of the centromeric sequences.

Our diploid genome assembly further revealed widespread variation in centromeric organization between the two haplotypes (Fig. S6). For instance, although the overall centromere lengths are comparable between the paternal and maternal copies of chromosome 14 (1.3 Mb vs. 1.4 Mb), the size of centromere cores differs by nearly two-fold (190 kb vs. 90 kb) (Fig. 2B). On chromosome 37, the relative position of the centromere core differed between the two haplotypes, suggesting either repositioning of the centromere core or differential expansion of pericentromeric satellite repeats (Fig. 2C). On chromosome 16, we detected a pericentric inversion that reversed the centromere organization (Fig. 2D; Fig. S7).

### Genetic variation of the W chromosome

The assembly of the avian W chromosome is challenging due to its heterochromatic and repetitive nature^34^. The assembled size of the W in GGswu2 is 23.5 Mb, approximately 26.9% of the size of the Z. Compared with the W chromosomes of GGswu1 and GRCg7b, ∼9 Mb and ∼12 Mb more sequences have been assembled, respectively (Fig. 3A). The PURs of the new chromosome W were mostly distributed in the intergenic regions, consisting of satellite DNA (Fig. 3B). The repetitive sequences accounted for 88.4% of the W, and satellite DNA was most abundant, accounting for 55.87% of all repeats. The tandem array of simple repeat (GGAGA)n has a size of ∼4.5 Mb, covering ∼19.3% of the W (Fig. 3B). A 21-bp element with dyad symmetry (AAAATnnnnnnTTTTCnnnnn) is the second most abundant repeat, with a total size of 4.2 Mb, and frequently formed divergent higher order repeats (HORs) (Fig. 3B).

**Fig. 3.**
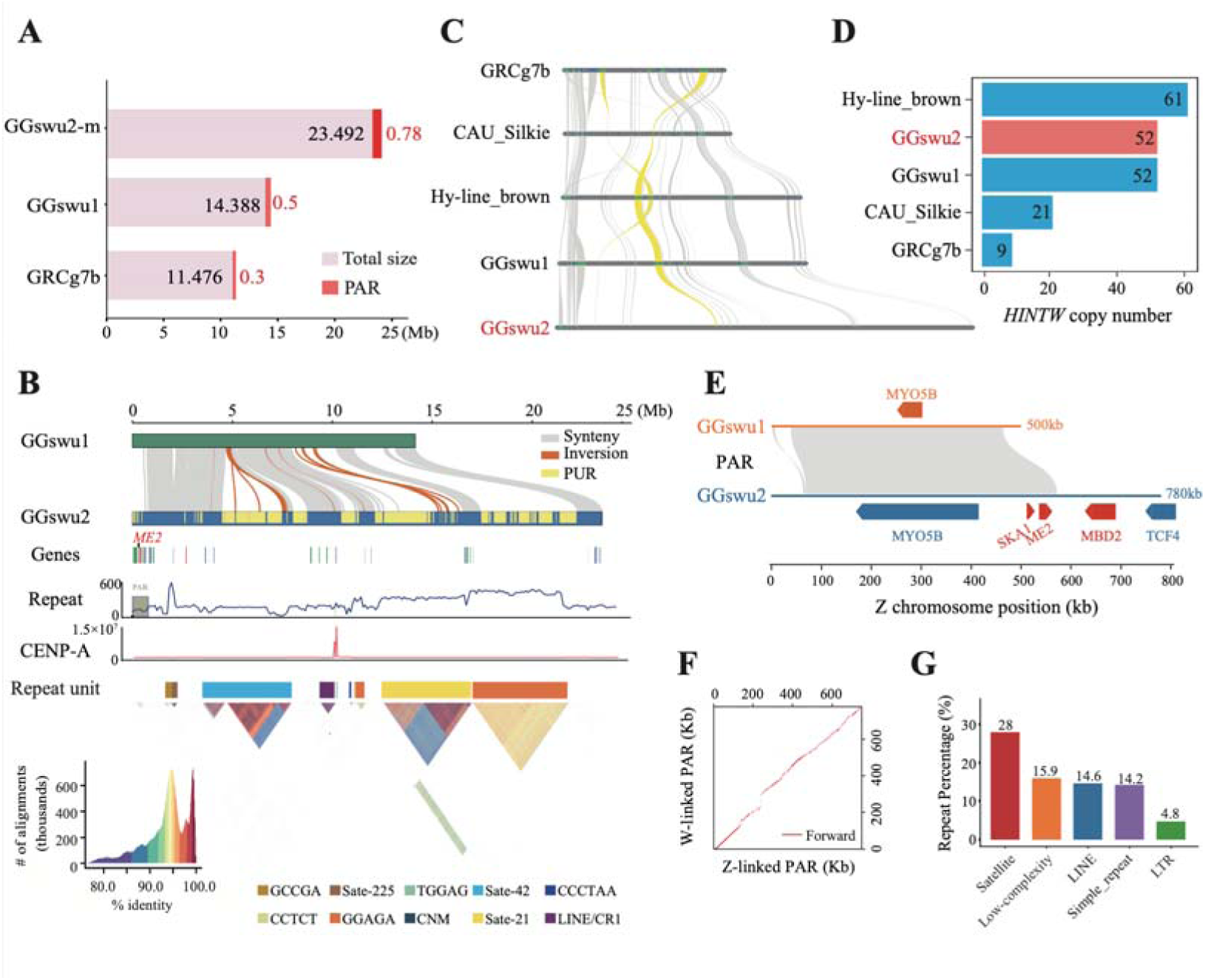
Improved assembly of the W chromosome. (**A**) Comparison of sizes of chromosome W and PAR among three chicken genomes. (**B**) The synteny and structure of the chromosome W. In the gene panel, the red vertical bars present newly identified genes, while the green and blue bars represent genes with positive and negative orientation, respectively. The heatmaps show the sequence identity (%) between 1 kb sequences. (**C**) The multi-collinearity analysis among several assemblies of chromosome W. The Hy-line_brown and CAU_Silkie breeds were obtained from two previous studies, respectively^13,52^. The yellow highlights indicate two representative gene rearrangement events. The violet and blue colors denote two copies of the *RPL17* gene. (**D**) Comparison of the copy number of *HINTW*. (**E**) PAR synteny between GGswu2 and GGswu1. The red colors represent the newly identified genes in the GGswu2. (**F**) The dotplot of PARs of the chromosome Z and W. (**G**) The repeat content of the PAR.

We further collected another four published high-quality chicken W chromosome assemblies and compare the distribution and copy number of protein-coding genes. All 28 conserved chicken W-linked genes reported by Bellott et al.^35^ were recovered in GGswu2, including *TCF4* whose gene structure remained incomplete in previous assemblies. The GGswu2 *TCF4W* gene model (484 amino acids) is more complete than the Bellott model (121 amino acids) when human and mouse homologs were used as reference; RNA-seq data from multiple tissues further supported our annotated gene model (Fig. S8). Overall, the gene order on the W chromosomes is conserved across different assemblies, though several genes, including *C18ORF25* and *NEDD4L*, were translocated (Fig. 3C). This is consistent with a recent cross-species analysis that demonstrated frequent inversions of bird W chromosomes^26^.

While most W-linked genes are single-copy, *HINT1W* is known to have multiple copies^34–37^, and we detected various copy numbers among the five W assemblies (Fig. 3D). Both GGswu1 and GGswu2 contained 52 copies of *HINT1W*, compared with 61 copies in Hy-line_brown, 21 copies in CAU_Silkie, and 9 copies in GRCg7b. Two copies of *RPL17* were previously reported in GGswu1 and are now confirmed in all other assemblies except for GRCg7b (Fig. 3D). In GGswu2, only one copy of *TXNL1* was detected, whereas three copies were identified in GGswu1. These differences in copy number among assemblies may reflect genuine copy-number variation, but differences in assembly completeness and repeat resolution, including potential repeat collapse, cannot be excluded.

Though the W chromosome has been highly differentiated from the Z, a part of it is still recombining with the Z, called the pseudoautosomal region (PAR). Our assembly recovered 0.78 Mb of PAR sequence, representing a 50% increase relative to GGswu1 (Fig. 3A). The assembled PAR contains three novel genes (Fig. 3B; Fig. 3E), including *ME2*, a gene that was thought to be missing in the chicken genome^26^. The Z- and W-linked PARs are highly conserved in synteny, but the difference in satellite DNA content may have contributed to the structural variation between them (Fig. 3F; Fig. 3G).

### Reduced nucleotide diversity of the dot chromosomes

Previous studies revealed distinct genomic features of the three types of the chicken genome: macrochromosomes, microchromosomes and dot chromosomes^21^. With the haplotype-resolved assembly of GGswu2, we set out to analyze the between-haplotype and within-species variations of chicken chromosomes. By aligning GGswu2-p and GGswu2-m, our analyses revealed conserved synteny between the two haplotypes for macro- and microchromosomes, but detected more frequent structural variations (SVs) for dot chromosomes (Fig. S9A). The enrichment of SNPs, INDELs and SVs on dot chromosomes was significant compared with both micro- and macrochromosomes (Fig. 4A). When we examined variant densities separately within genic/intergenic and repeat/non-repeat regions, the elevated INDEL and SV densities on dot chromosomes were largely retained in genic, intergenic and non-repeat regions; in contrast, SNP density differences among chromosome groups became statistically insignificant (Fig. S10).

**Fig. 4.**
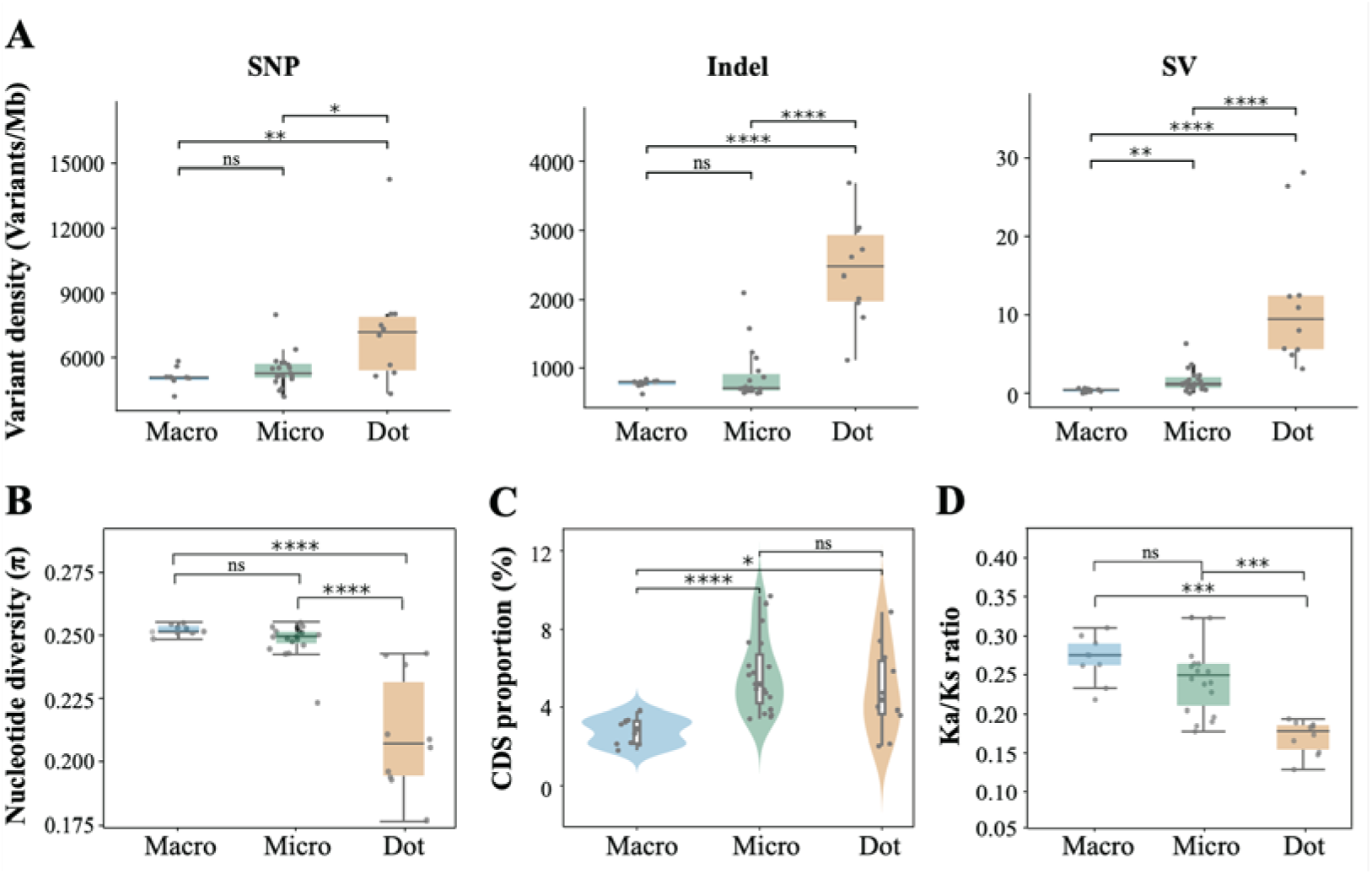
Genomic characteristics of dot chromosomes. (**A**) Chromosome-specific densities of SNPs, INDELs, and SVs. (**B**) Dot chromosomes have lower nucleotide diversity. (**C**) Microchromosomes and dot chromosomes have a higher density of CDS than that of macrochromosomes. (**D**) Dot chromosomes have the lowest Ka/Ks ratio. The Mann-Whitney U test was used to assess statistical significance. Statistical significance was indicated as follows: ns (not significant); * p<0.05; ** p<0.01; *** p<0.001; **** p<0.0001.

We then mapped resequencing data from 20 chicken populations to estimate nucleotide diversity (π) across chromosomes. Dot chromosomes exhibited much lower π than both macro- and microchromosomes (Fig. 4B; Fig. S9B). Stratification by genic/intergenic and repeat/non-repeat regions largely recapitulated the chromosome-level pattern (Fig. S11). Consistently, the dot chromosomes have a larger proportion of homozygous sites (Fig. S9C). This is likely because dot chromosomes have a higher density of coding sequences, which are the targets of purifying selection and background selection, leading to reducing nucleotide diversity^38^. Supporting this, our analyses revealed that the density of genes or coding sequences in dot chromosomes is higher than in macrochromosomes, though lower than in microchromosomes (Fig. 4C; Fig. S9D). Furthermore, our analyses of the Ka/Ks ratio showed the dot chromosomes significantly lower Ka/Ks ratio than that of microchromosomes, suggesting stronger purifying selection on the dot chromosomes (Fig. 4D). Together, our analyses suggested that the higher density of coding sequences, coupled with stronger purifying selection and background selection, have led to the lower π of the dot chromosomes.

### Dot chromosomes have a higher germline mutation rate

A recent pedigree-based estimate of germline mutation rate (GMR) of chicken using 3 individuals of one trio revealed a mutation rate of 3.63 × 10^−9 39^. In that study, GRCg6a was used as the reference that lacked considerable sequences of dot chromosomes and other chromosomes. To test whether the incomplete genome assembly impacts the estimate of GMR, we collected nine individuals that are siblings of the individual we used for the diploid genome assembly. Our estimated mutation rates varied from 2.43 × 10^−9^ to 9.57 × 10^−9^, with a mean value of 5.26 × 10^−9^, which was 1.4 times of the previous estimate^39^ (Fig. 5A; Table S8). We also detected on average 1.5 times more *de novo* mutations (DNMs) than the previous study. To assess whether the reference genome affected DNM detection and GMR estimation, we reanalyzed the same resequencing data against GRCg6a using the same pipeline. The mean genome-wide GMR estimated using GRCg6a (4.52 × 10^−9^) was 1.2-fold lower than that estimated using GGswu2, though higher than previous study^39^ (Table S9).

**Fig. 5.** The germline mutation rates of chicken chromosomes. (**A**) Germline mutation rates across macro-, micro-, and dot chromosomes in nine trios. Each point represents the estimated germline mutation rate for one chromosome type in an individual trio, and horizontal lines indicate the corresponding 95% confidence intervals (CIs). Different colors denote the three chromosome types. (**B**) Mean germline mutation rates across macro-, micro-, and dot chromosomes. Each grey spot represents one individual. The blue dashed line means the *de novo* mutation rate from Bergeron *et al*.^39^. (**C**) Feature-stratified germline mutation rates across chromosome types. Mutation rates were independently estimated for genic, intergenic, repeat, and non-repeat regions. Bars represent mean mutation rates across the nine trios, with error bars indicating standard errors (SEs).

The six dot chromosomes newly assembled in GGswu2 (chr29 and chr34-38), which are absent from GRCg6a, contained eight DNMs across the nine individuals (Table S10). These results indicate that the more complete GGswu2 assembly enabled detection of additional DNMs in previously missing regions. When calculated independently, the dot chromosomes showed the highest mutation rate, followed by microchromosomes. Specifically, the mutation rate of microchromosomes (5.08 × 10^−9^) was 1.4 times higher than that of macrochromosomes (3.67 × 10^−9^), while that of dot chromosomes (2.99 × 10^−8^) was about 8.1 times higher than macrochromosomes (Fig. 5B). To assess whether this pattern was associated with genomic composition, we further stratified mutation rates by genic and intergenic regions, as well as by repeat and non-repeat regions. The elevated mutation rate of dot chromosomes was also observed after stratification by genomic features (Fig. 5C). Dot chromosomes also showed substantially lower callable fractions and mapping qualities than macro- and microchromosomes across genomic features (Fig. S12), warranting cautions in interpreting their elevated mutation-rate estimates.

### Strong subtelomeric interactions of dot chromosomes

Unlike Hi-C data, the Pore-C data allowed us to capture longer and multi-way chromatin contacts^40,41^. We obtained 35.9 million reads which generated ∼0.41 billion pairwise interactions (Table S11), capturing more contacts by fewer reads compared to the Hi-C method^42^. We detected a positive correlation between the order of reads and contacts, consistent with the previous study of human^43^ (Fig. 6A). A concatemer could contain several contact fragments^41^, and from these fragments, we detected about 19 million concatemers and about 73% were high order (contact fragments >=3) (Fig. 6B). The most frequent multi-way contacts are 7- to 11-way (Fig. S13A), similar to the pattern reported in human genomes^43^. The chromosome 1 and 2 have most frequent multi-way contacts, likely because of their larger chromosome sizes (Table S12).

**Fig. 6.** The analysis of inter-chromosomal interactions based on Pore-C data. (**A**) The percentage of multi-way contacts. (**B**) The percentage of concatemers. (**C**) Mean interaction density of inter-chromosomal contacts among chromosome types. (**D**) Subtelomere-subtelomere contacts exhibit higher interaction density than other genomic region pairs. (**E**) Frequency and interaction density of subtelomere–subtelomere contacts among chromosome types. The Mann-Whitney U test was used to assess statistical significance. Statistical significance was indicated as follows: **** p<0.0001.

Our analysis of inter-chromosomal interactions detected strongest interactions between dot chromosomes (Fig. 6C), consistent with the previous analysis using Hi-C data^21^. We further found that the interaction between subtelomeric regions was significantly stronger than those involving other chromosomal regions (Fig. 6D). We then examined the frequency and interaction density of subtelomere-subtelomere contacts among different chromosome types. Dot-microchromosome pairs accounted for the highest number of subtelomeric contacts, indicating the greatest interaction frequency between these chromosome types. In contrast, dot-dot pairs showed the highest interaction density, suggesting that these contacts exhibited stronger contact signals despite contributing fewer contacts overall (Fig. 6E). Thus, chromosome pairs with the largest number of subtelomeric contacts were not necessarily those with the highest interaction density. By examining individual dot chromosomes, we further found that interactions between chr32 and chr34, chr35, and chr36 showed the highest interaction density among all chromosome pairs (Fig. S13B).

## Discussion

The pursuit of a complete and accurate reference genome is fundamental for deciphering the evolution and functionality of any species. Historically, the avian genome, exemplified by the domestic chicken (*Gallus gallus*), has been characterized as notably compact and simplistic among major vertebrate lineages^44^. This apparent simplicity was partly why the chicken was the first livestock genome sequenced in 2004 and led to the choice of birds for the first large-scale comparative genomic study^4^. Recent T2T and pan-genome analyses have since begun to dismantle the “simple genome” paradigm, revealing complexity hidden in repetitive subtelomeric regions, HORs of the centromeres, the large heterochromatic region of the dot chromosomes, frequent immune gene duplications (*e.g.* on chr16), and large tandem arrays of short repeats on the W chromosome^18,20,23,24,26,45^.

Our study generated a nearly complete, haplotype-resolved diploid assembly (GGswu2); however, at least 14 gaps remain unresolved, primarily within the short arms of the acrocentric chromosomes. The use of parental ONT sequencing data for trio-based haplotype phasing contributed to the improved haplotype resolution of dot chromosomes relative to GGswu1. Further improvements in ONT read accuracy and haplotype-resolved assembly algorithms^46,47^ are expected to facilitate more efficient and complete assembly of the short arms of these chromosomes.

Our results illuminate the unique evolutionary pressures acting on the dot chromosomes (chr16, 29-32, 34-38). These small chromosomes are recognized as hotspots of gene attrition, having lost approximately 29% of ohnologs surveyed, a rate far exceeding that of other avian chromosomes^22^. This massive gene loss suggests that the remaining genes are highly essential and subject to stronger purifying selection. Our analysis strongly supports this hypothesis: the dot chromosomes exhibit a significantly lower nucleotide diversity compared to macro- and microchromosomes. This reduced diversity, despite their size, is likely due to the highly gene-dense nature of these chromosomes acting as widespread targets for selective interference (such as background selection), which eliminates linked neutral variation^48^. Consistent with this strong selection, dot chromosomes show a significantly lower Ka/Ks ratio compared to microchromosomes. Paradoxically, simultaneously facing intense selective constraint, dot chromosomes also exhibit high instability. We found a much higher density of indels and SVs between haplotypes on dot chromosomes than on micro- or macrochromosomes, a pattern recently independently revealed in zebra finch^18^. This dynamic turnover is likely linked to their high GC content and prevalence of simple tandem repeats, which facilitate mechanisms like local repeat expansion or “sequence stuttering”^4,49^. This instability is thought to be driven by a high recombination rate coupled with active insertions of transposable elements (TEs), which mediate SV formation^50^. Beyond their distinctive evolutionary and sequence characteristics, our Pore-C analysis revealed enhanced interactions between subtelomeric regions. This pattern likely reflects the principles of avian nuclear chromosome organization where telomeric and subtelomeric repeats show non-random spatial distributions in interphase nuclei^51^, consistent with a recent study in zebra finch^18^.

### Limitations of the study

Although GGswu2 represents the most complete haplotype-resolved chicken genome assembly to date, residual gaps remain, and several gaps lack sufficient read coverage. While this study reveals inter-haplotypic variation in centromere sequences, the extent of such variation across populations remains to be determined. Furthermore, the completeness and accuracy of the W chromosome require further improvement to enable robust analysis of structural variation. We rely on short-reads to identify *de novo* mutations, but many pericentromeric and subtelomeric regions remain poorly accessible to short-read technologies. Consequently, the full spectrum of *de novo* mutations across the genome remains incompletely characterized.

## RESOURCE AVAILABILITY

### Lead contact

Requests for further resources should be directed to and will be fulfilled by the lead contact, Luohao Xu.

### Materials availability

This study did not generate new unique reagents.

### Data and code availability

- **Data:** The genome assemblies GGswu2-p and GGswu2-m were uploaded in NCBI under BioProject accession numbers PRJNA1467338 and PRJNA1467339, respectively. Raw sequencing data generated in this study, including PacBio HiFi data, ultralong ONT data, Pore-C data, whole-genome sequencing data, and RNA-seq data of the F1 chick; The ONT data and NGS data of parents, can be obtained from NCBI under BioProject accession number PRJNA1114433. The genome and annotation files are also deposited on Figshare (10.6084/m9.figshare.33191442).
- **Code:** This paper does not report original code.
- **Other:** Any additional information required to reanalyze the data reported in this paper is available from the lead contact upon request.

### Author contributions

L.X. and N.X. conceived and supervised the study. S.F. and L.X. wrote the manuscript. S.F, J.L. and N.K. performed bioinformatics analysis. D.L. and Z.L. contributed to the sampling. All authors read and approved the manuscript.

## Acknowledgments

This work was financially supported by the National Key Research and Development Program of China (2024YFF1001400), Open Project of the State Key Laboratory of Animal Biotech Breeding (2026SKLAB6-13) and the Chongqing Science Fund for Distinguished Young Scholars (CSTB2024NSCQ-JQX0021).

## STAR★Methods

### Key resources table

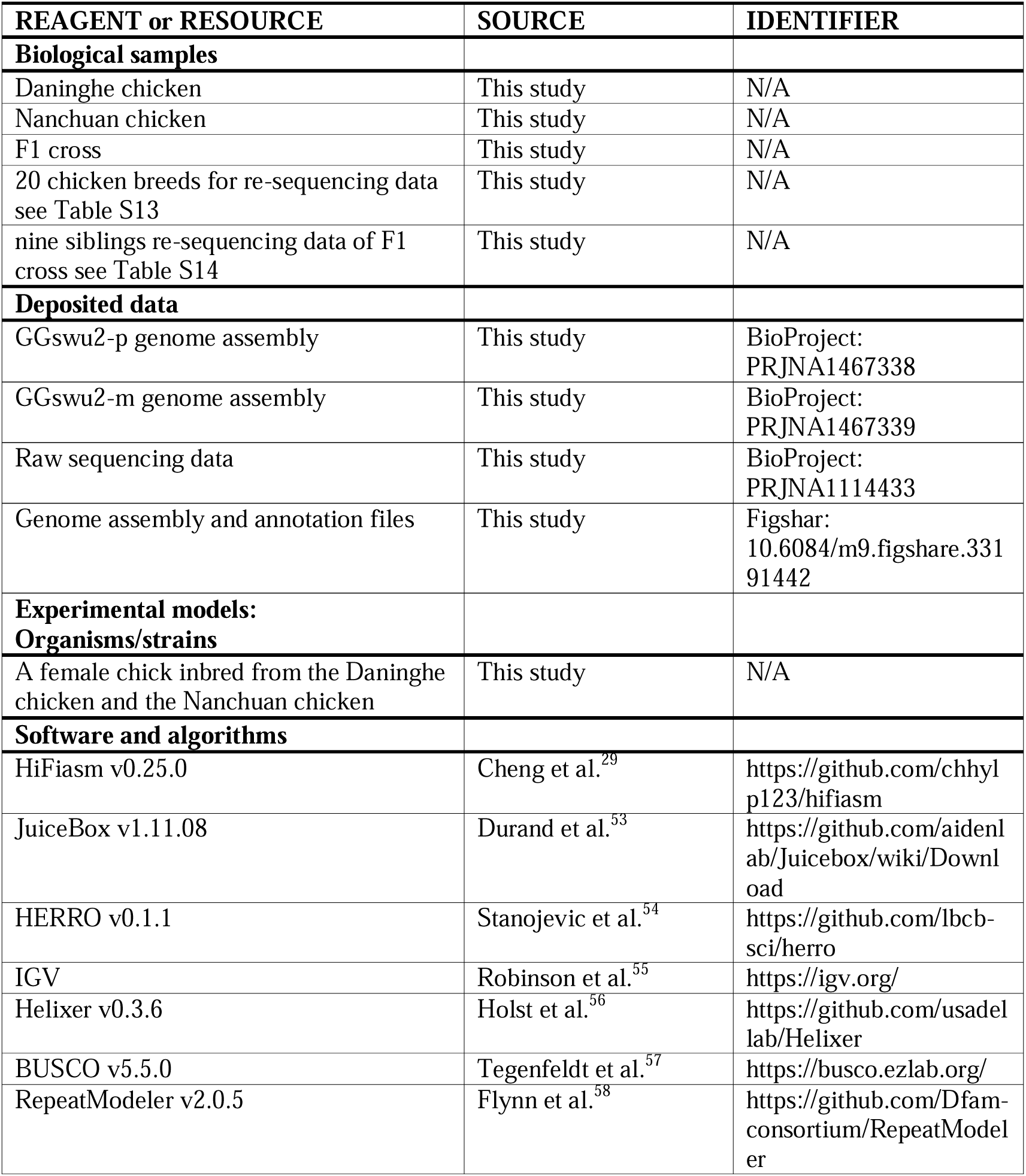
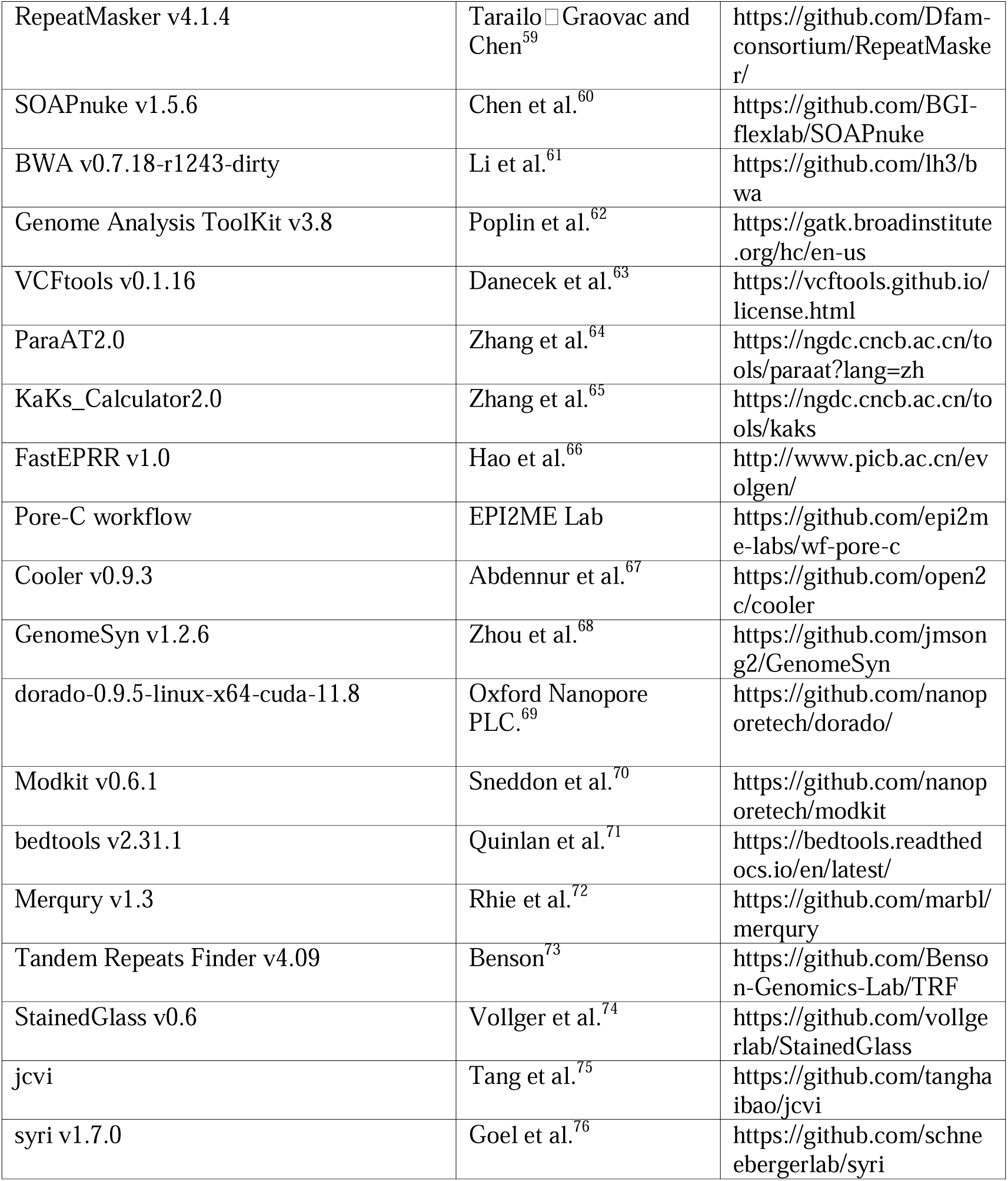

### EXPERIMENTAL MODEL AND STUDY PARTICIPANT DETAILS

A chicken trio was collected from Chongqing, China, comprising a maternal individual from the Nanchuan chicken breed, a paternal individual from the Daninghe chicken breed, and their F1 female offspring. Blood samples were collected from all three individuals for Oxford Nanopore Technologies (ONT) sequencing. Next-generation sequencing (NGS) was performed on both parents, whereas PacBio HiFi and Pore-C sequencing were performed on the F1 female offspring using blood-derived DNA. Total RNA was extracted from six tissues of the F1 female offspring, including the heart, spleen, lung, kidney, adipose tissue, and stomach, for transcriptome analysis.

### METHOD DETAILS

#### Genome Sequencing

We collected a chicken trio from Chongqing, with the maternal individual from the Nanchuan chicken breed and the paternal individual from the Daninghe chicken breed. The parents and F1 female chick were all sampled for Oxford Nanopore Sequencing (ONT), and we also did the Next-Generation Sequencing (NGS) for the parents, and PacBio HiFi sequencing and Pore-C sequencing for the F1 chick. For each ultralong Nanopore library, approximately 8 to 10 μg gDNA was size-selected (>50 kb) with SageHLS HMW library system (Sage Science) and processed using the Ligation sequencing 1D kit (SQK-LSK109, Oxford Nanopore Technologies). About 800 ng DNA libraries were constructed and sequenced on the Promethion platform at Grandomics Biosciences (Wuhan, China). For HiFi (CCS) sequencing, SMRTbell target size libraries were constructed according to PacBio’s standard protocol (Pacific Biosciences) using the 15-kb preparation solutions. A total amount of 15 μg DNA per blood sample was used for the DNA library preparations. Sequencing was performed on a PacBio Sequel II instrument with Sequencing Primer V2 and Sequel II Binding Kit 2.0 at Grandomics. For Pore-C sequencing, 1 μg of genomic DNA input was used to generate a sequencing library according to the protocol provided for the SQK-LSK109 kit (Oxford Nanopore Technologies, Oxford Science Park, UK, version GDE_9063_v109_revU_14Aug2019). The sample was wrapped with gauze and immersed in cross-linking solution containing 1% (v/v) formaldehyde under vacuum for 30-40min. The glycine was added to the solution to quench the cross-linking reaction. The sample was washed with pre-chilled ddH_2_O and ground into powder in liquid nitrogen. Then the sample blended by overturn after the addition of HB buffer. Nuclei were extracted by centrifuging the filtrate at 2500g. DNA manipulation was performed after in situ digestion using the DpnII restriction endonuclease, in situ proximity ligation, reverse-crosslinking process. Pore-C library was finally sequenced on the PromethION sequence.

#### RNA Sequencing

Total RNA was extracted from six tissues (heart, spleen, lung, kidney, fat, and stomach) using TRIzol reagent (TIANGEN) according to the manufacturer’s instructions. RNA libraries were constructed from 1 μg of total RNA using the TruSeq RNA Library Preparation Kit (Illumina, USA). Poly(A)+ mRNA was enriched with oligo(dT) magnetic beads and converted into cDNA following the manufacturer’s protocol. After adapter ligation and PCR amplification, the libraries were purified, quality-checked using an Agilent Bioanalyzer 2100 (Agilent Technologies, USA), and sequenced on the Illumina NovaSeq platform to generate 150 bp paired-end reads.

#### Genome Assembly and gap-filling

Trio-binning was used to assemble the haplotype-resolved genomes based on HiFi, ONT and the parents’ sequencing reads, using HiFiasm (v0.25.0)^29^. The Pore-C data was used to facilitate scaffolding of the contigs and manual curation was performed in JuiceBox (v1.11.08)^53,77^.

To fill the gaps, ONT reads were corrected using HERRO (v0.1.1)^54^ with standard parameters. We then used the corrected ONT data as input for HiFiasm assembly, again with the trio mode. The generated haplotype-resolved contigs were used for gap-filling of the corresponding haplotype assemblies. When a contig was aligned to both flanking sequences of a gap, it was considered to span the entire gap, and the gap region was replaced with the aligned sequence from the contig. This process filled 52 gaps in GGswu2-p and 43 gaps in GGswu2-m. Secondly, we combined HiFi and corrected ONT data for the HiFiasm assembly, with similar parameters. The generated contigs were used for the second round of gap-filling, resolving 9 gaps in GGswu2-p and 7 gaps in GGswu2-m.

To systematically assess the sequence continuity and read support of the filled regions, we evaluated each gap using independent ONT and HiFi reads. Mean read coverage was calculated separately for each filled region and for the corresponding remainder of the chromosome. Full-length spanning reads were defined as reads whose alignments extended beyond both boundaries of the filled region. For longer filled regions where individual reads did not span the entire sequence, continuous support was assessed by merging the alignment intervals of primary reads overlapping the filled region. Overlapping or directly adjacent alignment intervals were merged, and any uncovered intervals between merged blocks were recorded as breakpoint regions. A filled region was considered continuously supported when the merged read alignments covered the entire filled region without any breakpoint. Detailed per-gap coverage and read-support statistics are provided in Table S2-S3.

Representative filled regions were further examined using IGV^55^. For each region, the filled sequence and 100-bp flanking regions on both sides were visualized to assess read support across both junctions and throughout the inserted sequence (Fig. S1).

#### Genome Annotation

Gene prediction was carried out using Helixer (v0.3.6)^56^, a deep learning-based annotation tool, with the soft-masked genome as input. Helixer was executed within a Singularity container using the vertebrate lineage model and corresponding pretrained weights^56^. Subsequently, we detect GFF3 format error using gff3_QC and automatically correct them using gff3_fix (https://github.com/NAL-i5K/GFF3toolkit). We assessed the completeness and overall quality of the annotated gene set using BUSCO (v5.5.0)^57^ based on conserved orthologs. Repeats were analyzed via a method that combines *de novo* structure analyses and homology comparisons. First, RepeatModeler (v2.0.5) ^58^was employed to construct the repeat element library. Then, the repeat library was then used to perform repeat annotation using RepeatMasker (v4.1.4)^59^.

#### Genome evaluation

We adopted multiple indicators to evaluate the completeness and accuracy of the *de novo* assemblies. First, we calculated the mapping rates as an evaluation metric. Six high-quality chicken resequencing datasets were downloaded from NCBI and aligned to different versions of the chicken reference genome using default parameters of BWA-MEM^61^. The average mapping rates were then calculated (Table S15). Next, we used BUSCO scores to assess assembly completeness^57^. CRAQ, a reference-free genome assembly evaluator, can assess the accuracy of assembled sequences and provide a detailed multi-perspective quality assessment. It reports overall assembly quality index (AQI) metrics, including regional and structural assembly quality (R-AQI and S-AQI)^30^. Therefore, we used HiFi data to compute the AQI for all reference genomes and took the average of the sum of R-AQI and S-AQI as the final result (Table S16). Additionally, we used Merqury to evaluate the assemblies by comparing k-mers derived from unassembled, high-accuracy sequencing reads with the genome assembly, which provides estimates of genome consensus quality (QV) and haplotype completeness^78^. An overall k-mer evaluation was performed using k-mer multiplicities from combined long reads (ONT and HiFi). The spectra-asm plot illustrates shared and haplotype-specific k-mers, reflecting diploid assembly completeness by showing the fraction of k-mers unique to each of the primary and alternate assemblies (Fig. S14). Finally, we used GCI, another assembly assessment tool for high-quality genomes, which reports a score quantifying assembly continuity^31^. As the developer previously benchmarked GGswu1 using both HiFi and ONT data, we also calculated GCI for the new assemblies based on the same data types.

#### Methylation calling

We used dorado (dorado-0.9.5-linux-x64-cuda-11.8)^69^ to detect 5-methylcytosine (5mC). Subsequently, Modkit (v0.6.1) ^70^was employed to rapidly convert the resulting modBAM files into bedMethyl format. Finally, bedtools (v2.31.1)^71^ was used to summarize the methylation frequency across 50 kb windows.

#### Nucleotide diversity

We collected NGS data from 20 chicken breeds originating from diverse geographical regions both domestically and internationally (Additional Data S6). Raw reads were trimmed by SOAPnuke (v1.5.6)^60^ software to filter out low-quality bases and sequences. High-quality reads were aligned against the reference genome (GGswu2-p merged with chromosome W of GGswu2-m) using bwa “BWA-MEM” algorithm. Alignment files were next subjected to a series of processing and filtrations including position sorting, duplicated read marking and removal, which were carried out using Picard (v1.56)^79^. SNPs were called using the HaplotypeCaller and GenotypeGVCFs commands in the Genome Analysis ToolKit (v4.2.6.1)^80^, then filtered using the VariantFiltration with the following parameters: QD < 2.0, FS > 20.0, MQ < 40.0, MQRankSum < –2.0, MQRankSum > 4.0, ReadPosRankSum < –3.0, ReadPosRankSum > 3.0, and SOR > 3.0. VCFtools (v0.1.16)^63^ was used to calculate the nucleotide diversity (π) of the different chromosomes. We used parameter “-- site-pi” to calculate π in every site of chromosomes and “-- window-pi 50000 -- window-pi-step 25000” to set the window and step when divided by windows.

#### Synonymous mutations and nonsynonymous mutations

We firstly collected the genomic data of the recently released mallard genome^52^, and sought out the orthologs of chicken (GGswu2-p) and mallard using Orthofinder software (v2.5.5)^81^. Then, we aligned protein sequence of orthologs in parallel and translated them into DNA sequence by ParaAT2.0^64^. Finally, we used KaKs_Calculator2.0^65^ to calculate Ks, Ka and their ratio.

#### The calculation of recombination

We did the whole-genome sequencing for 9 siblings of the chick that was used to reference genome assembly (Additional Data S7) and applied the same method as the above to call variants and processed filters. After obtaining the final combined VCF file, we used FastEPRR (v1.0)^82^ to calculate the rate of recombination of every chromosome (window size 100 kb).

#### Germline mutation rates

As above, we collected data of 9 siblings of the F1 chick to estimate the germline mutation rates (GMR; Table S14) and applied a pedigree-based analysis pipeline of DNM^83^. Raw reads were trimmed with SOAPnuke (v1.5.6)^60^. The mapping was conducted with BWA-MEM (v0.7.18)^61^. We used the maternal haplotype GGswu2-m as a reference genome. A post-mapping step removed any reads mapping to multiple regions of the genome as well as duplicated reads using Picard MarkDuplicates (v2.27.4)^79^. We called variants for each individual using HaplotypeCaller in BP-RESOLUTION mode with GATK (v4.2.6.1)^80^. This mode returns a genotype quality and depth for all positions of the genome, not only the polymorphic sites. As recommended by GATK best practices, GenomicsDBImport combined all gVCF files per species into a single file and GenotypeGVCF applied a joint genotyping of all samples. Similar filtering methods to those in the previous study were then applied to detect DNMs^39,83^. Therefore, for each trio, we applied the following filters:

(1) We used a site filtering with the following parameters: QD < 2.0, FS > 20.0, MQ < 40.0, MQRankSum < –2.0, MQRankSum > 4.0, ReadPosRankSum < –3.0, ReadPosRankSum > 3.0, and SOR > 3.0.
(2) For Mendelian violations, variants that deviated from Mendelian inheritance were selected using GATK SelectVariant and refined with an R script to keep only sites in which both parents were homozygous for the reference allele (HomRef), and the offspring was heterozygous (Het).
(3) For depth filter (DP), only positions with a DP > 0.5 × m_depth_ and DP < 2 × m_depth_ for each individual were kept, with m_depth_ being the average depth of the trio.
(4) Our allelic balance filter cut-off was 30–70% of the reads supporting the alternative allele, similar to previous studies^39,83^.
(5) For genotype quality filter (GQ), to ensure that only high-quality genotypes were retained for the analysis of trios, we removed all sites where one individual of the trio had a GQ <60.

We then calculated the number of callable sites, *C*, i.e., the number of sites in the genome where we would be able to call a *de novo* mutation if it was there for each trio. Specifically, based on the unique bam after a series of post process, we have selected high-coverage, high-quality callable sites, with filtering parameters set as: DP>=10, and MAPQ>=20. It is necessary to correct the false-negative phenomenon. We estimated the false-negative rate *α*, defined as the proportion of true heterozygote sites (1 parent: HomRef, another: HomAlt, and their offspring: Het) outside the allelic balance threshold. We also implemented in this parameter the false-negative rate of the site filters following a normal distribution (FS, MQRankSum, and ReadPosRankSum). In addition, we manually checked the read mapping quality for all *de novo* mutation sites in IGV^55^ (Fig. S14), and we removed the false-positive calls for which the variant was absent from the offspring or also present in a parent (Table S17). Finally, combining these parameters, the mutation rate per site per generation can then be estimated per trio with the following equation: GMR = DNMs / ((1 – *α*) × 2 × *C*).

To assess whether differences in genomic composition, callability, or mapping quality contributed to variation in mutation rates among chromosome types, we further stratified the analysis by genomic features. Genic and intergenic regions were defined based on genome annotations, while repeat and non-repeat regions were defined using RepeatMasker annotations. For each trio, mutation rates were calculated separately for macro-, micro-, and dot chromosomes within each genomic feature using the corresponding callable sites as the denominator. Callable regions were defined using a minimum read depth of 10 and a minimum mapping quality (MAPQ) of 20, following the criteria used for the genome-wide mutation-rate analysis. The callable fraction was calculated as the proportion of bases within each chromosome type and genomic feature that met these criteria. To further evaluate mapping and coverage characteristics within callable regions, we quantified the distribution of MAPQ and read depth across genic, intergenic, repeat, and non-repeat regions for each chromosome type.

#### Pore-C data analysis

We obeyed a Pore-C workflow for deriving high-order chromatin contacts from Pore-C concatemers (https://github.com/epi2me-labs/wf-pore-c). Briefly, the workflow involves alignment of Pore-C reads to a reference genome using Minimap2 (v2.28)^84^. The resulting alignments are filtered to identify a minimal alignment set that covers each Pore-C read. The Pore-C python package is then used again to filter spurious alignments, detect ligation junctions and assign chimeric fragments. The aligned segments will be used to generate a “walk” which enumerates the alignment coordinates of the monomers comprising the chimeric read and this is used to annotate the alignments. The pipeline will output the tagged alignments in a name sorted and coordinate sorted BAM. In additions, it also can output pairs and cooler formats for downstream analysis. The output cooler file contained the information of contact matrix, and we used Cooler (v0.9.3)^67^ to read it and did downstream analysis based on it.

### QUANTIFICATION AND STATISTICAL ANALYSIS

Statistical analyses were performed using R (v4.4.1). The Mann–Whitney U test was used to assess statistical significance, as indicated in the corresponding figure legends. Details of the statistical analyses, including sample sizes (n) and the definition of n, are provided in the corresponding figure legends and Results section. Statistical significance was indicated as follows: *p* < 0.05, p < 0.01, *p* < 0.001, and p < 0.0001.

## Supplemental information titles and legends

**Document S1. Figure S1-S15 and Table S1, Table S4, Table S6, Table S9-S12, and Table S15-S16.**

**Figure S1.** IGV visualization of representative gap-filled regions from the two GGswu2 haplotypes. Representative gap-filled regions on GGswu2-p: Chr32 and GGswu2-m: Chr1 are shown, with 100 bp of flanking sequence on each side. ONT and HiFi read alignments are displayed to assess read support across the filled regions and their junctions.

**Figure S2.** The re-sequencing data mapping rate and novel gene distribution. (**A**) The average mapping rate based on six high-quality re-sequencing data of chicken samples from NCBI. (**B**) The distribution of novel genes which were absent from GGswu1(Ren et al.)

**Figure S3.** The heatmap of sequence and epigenetic features across all chromosomes in GGswu2-m.

**Figure S4.** The altered centromeres in GGswu2-p.

**Figure S5**. The altered centromeres in GGswu2-m. chromosome 15 and chromosome 37 of GGswu2-m are shown in Fig. 2.

**Figure S6**. Variation in centromeric organisation between two haplotypes.

**Figure S7**. A pericentric inversion occurred on chromosome 16, with the inversion breakpoints supported by HiFi reads visualized in IGV.

**Figure S8.** RNA and Protein-level evidence supporting the completeness of *TCF4* gene model in GGswu2. (**A**) RNA-seq evidence supporting the *TCF4* gene model on the GGswu2 W chromosome. RNA-seq reads from three GGswu2 tissues (kidney, liver, and lung) were merged, mapped to the GGswu2 W chromosome, and visualized using IGV. (**B**) Protein sequence evidence supporting the *TCF4* gene model on the GGswu2 W chromosome. Multiple sequence alignment of *TCF4* proteins from Bellott, GGswu2, human, and mouse.

**Figure S9.** Genomic characteristics of dot chromosomes. (**A**) The collinearity analysis between two haploid genomes. The top one is GGswu2-p, and bottom is GGswu2-m. (**B**) The nucleotide diversity in the single sites of every chromosome. (**C)** The genotypes of different chromosomes. (**D**) Comparison of gene densities of the different types of chromosomes.

**Figure S10.** Variant densities across chromosome groups stratified by genomic features.

**Figure S11.** Nucleotide diversity across chromosome groups stratified by genomic features.

**Figure S12.** Callable fractions and mapping quality assessment across chromosome types and genomic features. (**A**) Callable fractions across chromosome types and genomic features. Bars represent mean callable fractions across the nine trios, with error bars indicating SEs. (**B**) Mapping quality assessment across chromosome types and genomic features.

**Figure S13.** The rule of inter-chromosomal interaction. (**A**) The multi-way contact patterns of chicken and human. (**B**) Top 20 chromosome pairs involved in subtelomere-subtelomere interactions. left: top 20 pairs by interaction frequency; right: top20 pairs by interaction density.

**Figure S14**. Merqury plots for evaluating k-mer completeness and haplotype phasing. (**A**) Distinct k-mer assembly spectrum (spectra-asm) plot for evaluating k-mer completeness. (**B**) Hap-mer blob plot for evaluating haplotype phasing.

**Figure S15.** The false-positive sites of germline mutation rate. Left: c1 (offspring), chr3: 93962203; Right: c3 (offspring), chr29: 590254. cp: father; cm: mother.

**Table S1**. Basic statistics of ultralong ONT reads.

**Table S4**. Coordinates of unfilled gaps in the GGswu2 genome.

**Table S6**. Sequence characteristics of GGswu2 chromosomes (GGswu2-m).

**Table S8**. Summary of germline mutation rate analyses.

**Table S9**. Comparison of GMR between GRCg6a and GGswu2 based on the same pedigree data.

**Table S10**. *De novo* mutations detected on the six dot chromosomes newly assembled in GGswu2

**Table S11.** Statistics of Pore-C reads

**Table S12.** The top inter-chromosomal contacts of the 7-11-way contacts

**Table S14.** Nine siblings re-sequencing data of F1 cross.

**Table S15**. Sample information for genome mapping rate analysis.

**Table S16.** Statistics of R-AQI and S-AQI across different genomes

**Data S1. Table S2-S3, Table S5, Table S7, Table S13, and Table S17.**

**Table S2.** Gap-filling validation data for GGswu2-p.

**Table S3.** Gap-filling validation data for GGswu2-m.

**Table S5.** PURs in every chromosome of GGswu2.

**Table S7**. PURs in short arms of acrocentric chromosomes.

**Table S13**. 20 chicken breed samples subjected to whole-genome resequencing.

**Table S17**. False-positive DNMs on chromosomes.

